# Recent thymic emigrants are preferentially recruited into the memory pool during persistent infection

**DOI:** 10.1101/2025.02.06.636722

**Authors:** Zachary T. Hilt, Arnold Reynaldi, Megan Steinhilber, Shide Zhang, Samantha P. Wesnak, Norah L. Smith, Miles P Davenport, Brian D. Rudd

**Author notes:** To whom correspondence should be addressed: Dr. Brian D. Rudd, Department of Microbiology and Immunology, C5 147 VMC, Cornell University, Ithaca, NY 14853.

## Abstract

Cytomegalovirus (CMV) leads to a unique phenomenon known as ‘memory inflation,’ where antigen-specific memory CD8+ T cells continue to accumulate in the peripheral tissues during the latent stage of infection. However, it is still not clear how the inflating pool of memory CD8+ T cells is generated and maintained. In this study, we used murine cytomegalovirus (MCMV) as a model of persistent infection and fate-mapping mice to determine the dynamics of CD8+ T cell recruitment into the memory pool. We found that neonatal exposure to CMV leads to an expansion of newly made CD8+ T cells (recent thymic emigrants, RTEs), which are maintained in the long-lived memory compartment. In contrast, CD8+ T cells made during the latent phase of infection (mature CD8+ T cells) contribute little to the memory pool. We also observed notable phenotypic differences between RTEs and mature cells. Whereas RTEs present at the time of infection gave rise to more effector memory cells, the cells produced later in infection were biased towards becoming central memory cells. Importantly, the preferential recruitment of RTEs into the effector memory pool also occurs during adult exposure to CMV. Collectively, these data demonstrate that persistent infection expands the RTE population, and timing of infection dictates whether neonatal or adult RTEs are ‘locked in’ to the memory pool.

**Author Summary:** Following infection with CMV, CD8+ T cells accumulate in the blood and peripheral organs over time, a feature termed ‘memory inflation’. However, it is not clear whether memory inflation is due to the continuous recruitment of cells made during the latent stage of infection or expansion of CD8+ T cells that were present at the time of infection. To address this question, we used a fate-mapping mouse model and examined the recruitment of CD8+ T cells that were produced during different stages of infection. Surprisingly, we discovered that CD8+ T cells exported from the thymus just prior to infection are preferentially recruited and maintained in the memory pool. In contrast, CD8+ T cells made during the latent stage contribute minimally to the inflating pool and exhibit a less differentiated phenotype. These results provide a new conceptual framework for understanding how the memory pool is generated and maintained after persistent viral infection.

## Introduction

Cytomegalovirus (CMV) is one of the most ubiquitous persistent viral infections in humans worldwide [1]. Depending on the geographical location, 12-56% of people become seropositive for CMV within the first year of life [2]. Infection with CMV leads to a phenomenon known as ‘memory inflation,’ where antigen-specific CD8+ T cells accrue over time [3–8]. These inflating CD8+ T cells have an effector memory phenotype characterized by low expression of CCR7, CD62L, CD27, and CD127 and high expression of KLRG1 and CX3CR1 [4, 9]. Despite being terminally differentiated, CMV-specific cells do not express markers of exhaustion (e.g., PD1, Lag3) and remain functional throughout the course of infection [4, 10]. However, a major issue is that CMV-specific CD8+ T cells continue to accumulate with progressing age and can occupy up to 50% of the total memory CD8+ T compartment [11]. In humans, expansion of CMV-specific cells is associated with higher all-cause mortality in older adults [12, 13].

A prevailing notion in the field is that the accumulation of CMV-specific T cells corresponds to the priming of naïve and/or memory CD8+ T cells that occurs during viral reactivation events [4, 14, 15]. However, recent work has shown that after initial infection with CMV, the virus does not need to replicate or spread from cell to cell to generate memory inflation [16]. Moreover, Loewendorf et al. demonstrated that memory inflation is maintained in mice after thymectomy, suggesting that recruitment of cells produced after infection is not necessary [15]. Thus, it remains unclear how the inflationary pool of CD8+ T cells is generated and maintained throughout the course of infection.

In the past, T cell immunologists have relied on adoptive cell transfers to determine which subsets of CD8+ T cells are recruited into the memory pool after infection [17, 18]. Alternative methods to measure cell recruitment include immune depletion and reconstitution, such as busulfan treatment of recipient mice, to facilitate bone marrow transplantation without killing established memory T cells [4, 19]. However, we reasoned we could obtain new insight into the underlying dynamics of memory inflation by leveraging a fate-mapping system that allows us to track waves of CD8+ T cells produced in the thymus at different periods of life without perturbing immune development or the ongoing infection [20–22]. This strategy exploits a TCRδ promoter-driven, tamoxifen-inducible cre (TCRδ-creERT2) to permanently mark, or ‘timestamp,’ CD8+ T cells that are present in the thymus at the time of tamoxifen exposure. The labeled cells can then be tracked in the periphery for the life of the animal. The advantage of this approach is that we can directly measure the recruitment of endogenous CD8+ T cells after CMV infection with minimal manipulation of the animals. Here, we use our timestamp mice to elucidate how memory inflation is maintained in mice infected with CMV at different stages of life.

## Results

### Neonatal exposure to CMV drives memory inflation of CD8+ T cells

Since CMV infection is common in early life, we felt it was important to study memory inflation in younger animals. However, less is known about the process of memory inflation after neonatal infection, so we first sought to determine how the numbers and phenotype of antigen-specific CD8+ T cells change after MCMV infection in early life. For these studies, we used a recombinant strain of MCMV that expresses the HSV-1 gB-8p peptide (MCMV-gB) that leads to an overwhelmingly dominant gB-specific CD8+ T cell response [23, 24], allowing us to monitor the antigen-specific CD8+ T cell response by tracking a single epitope. Mice were infected with MCMV-gB at birth to model pediatric infections, while uninfected control mice were given PBS (Fig 1A). CD8+ T cells were collected from the spleen at 1, 2, 4, 8, 12 and 21 weeks of age and stained with antibodies and tetramer (Kb:gB^498-505^) to phenotype the cells at each timepoint. We found that antigen-experienced CD8+ T cells (CD49d+CD44+) accumulated within the CD8+ T cell compartment, and this accumulation began to level off at around ∼60% of total CD8+ T cells by >5 weeks post infection (S1A Fig). We also measured the number of tetramer positive cells and found that ∼20% of CD8+ T cells were gB-specific at 12 weeks of age (Fig 1B). Thus, newborns infected with MCMV mount a robust memory inflation response.

**Figure 1.**
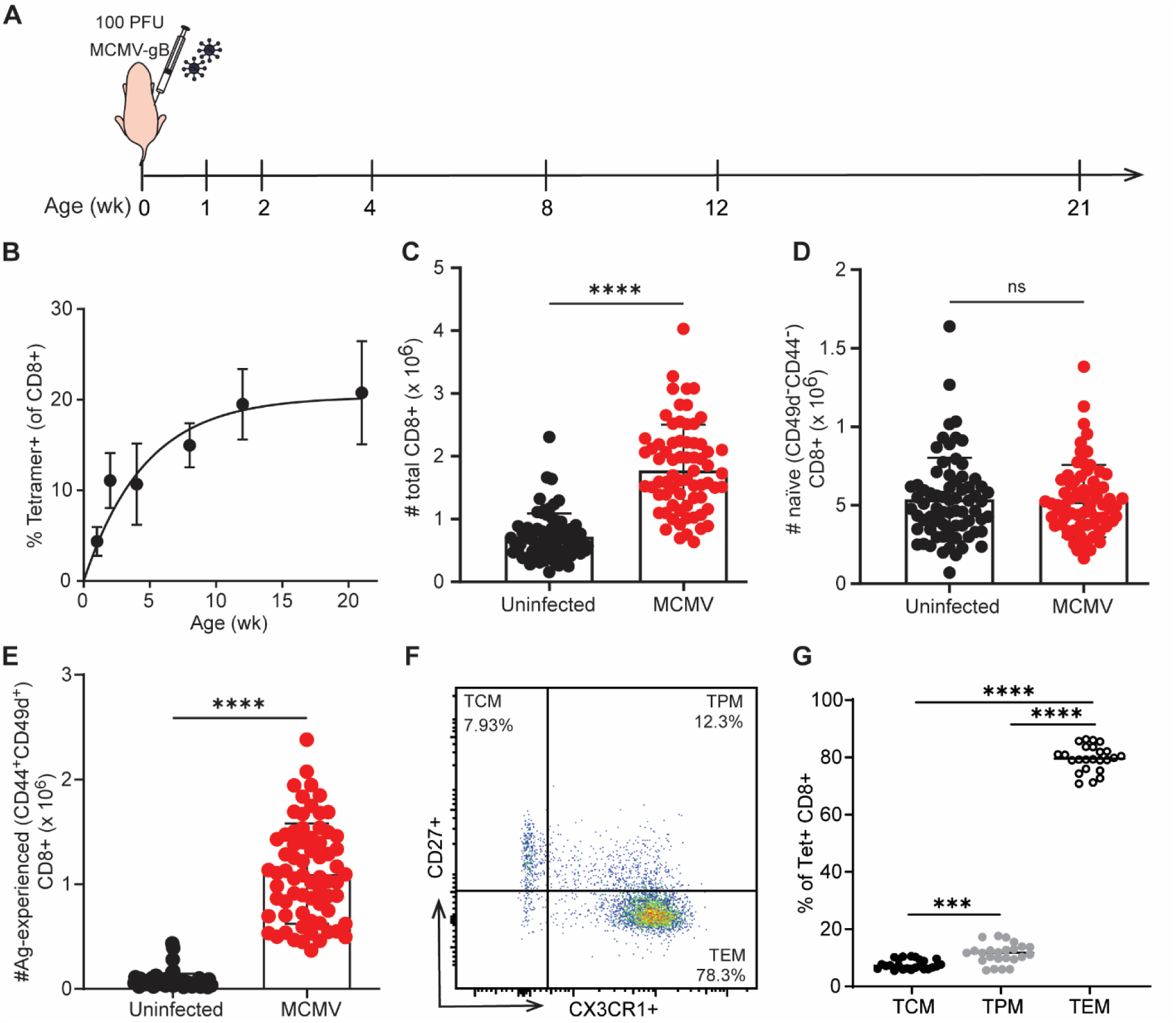
Neonatal exposure to CMV drives a robust memory inflation response. (A) Schematic of experimental design. Newborn mice were infected with MCMV-gB at birth. CD8+ T cells were isolated from the spleen at 1, 2, 4, 8, 12 and 21 weeks post-birth. (B) CD8+ T cells were stained with gB-tetramer specific cells and examined by flow cytometry (N = 4-10 mice). At 21 weeks, the total number of (C) CD8+ T cells, (D) naïve (CD49d-CD44-) and (E) antigen-experienced (CD44+ CD49d+) CD8+ T cells were quantified. Within the tetramer sub gates memory CD8+ T cells were identified as Central Memory (TCM, CD27+ CX3CR1-), Peripheral Memory (TPM, CD27+ CX3CR1+) or Effector Memory (TEM, CD27-CX3CR1+). (F) Representative dot plots of the gating scheme and (G) quantification of each memory subset are represented. For statistical test of two-groups an unpaired t-test with Mann-Whitney test for correction was performed. For statistical test of more then two-groups, an ordinary One-way ANOVA with Tukey’s multiple comparisons test was performed. Results are shown as mean ± SD or mean only. ***p<0.001, ****p<0.0001.

We next wanted to know if the expansion of memory cells in neonatal-infected mice impacts the numbers of naïve CD8+ T cells (CD49d-CD44-). To test this, we bled another group of mice and examined the number and phenotype of CD8+ T cells at 16 wks of age. We observed an increase in the total number of cells in the bulk CD8+ T cell compartment (Fig 1C). However, neonatal infection with MCMV did not alter the number of naïve CD8+ T cells. Instead, the observed increase was due to the significantly larger numbers of antigen-experienced CD8+ T cells (Fig 1D-E). These findings are consistent with results obtained in adult MCMV infection models, showing the large expansion of antigen-experienced CD8+ T cells after MCMV does not come at the expense of the naïve compartment [25, 26].

We next wanted to determine the phenotype of the inflating memory CD8+ T cells. Memory CD8+ T cells can be classified as central memory (CX3CR1-CD27+), peripheral memory (CX3CR1+ CD27+), or effector memory (CX3CR1+ CD27-) [27, 28]. At 21 weeks post infection, the antigen-experienced CD8+ T cell pool was comprised of approximately ∼60% central memory, ∼30% effector memory, and ∼8% peripheral memory (S1B-C Fig). In contrast, the tetramer positive CD8+ T cells were made up of approximately ∼8% central memory, ∼80% effector memory, and ∼12% peripheral memory (Fig 1F-G). These numbers are consistent with previous reports showing that memory inflation during CMV infection as an adult leads to an expansion of antigen-specific CD8+ T cells with a terminally differentiated phenotype [4]. Collectively, our data demonstrates that neonatal infection elicits an inflating memory pool that exhibits many of the key features observed in adult infection.

### Neonatal memory inflation is comprised of cells made closest to the time of infection

An important question is whether this sustained memory inflation response in neonates was maintained by a self-renewing pool or recruitment of newly made CD8+ T cells during adulthood. To address this question, we used our fate-mapping mouse model to mark the CD8+ T cells that are produced at different times after infection. Since CD8+ T cells made in early life are poor at forming memory, it is possible that the inflating pool is sustained by cells made later in life [29]. On the other hand, neonatal CD8+ T cells persist into adulthood and respond more quickly than adult-derived CD8+ T cells, which may enable them to preferentially respond to reactivating antigens during the latent phase [22, 29]. To distinguish between these possibilities, we administered tamoxifen at 1, 7, and 28 days post birth to ‘mark’ different developmental layers of CD8+ T cells (Fig 2A).

**Figure 2.**
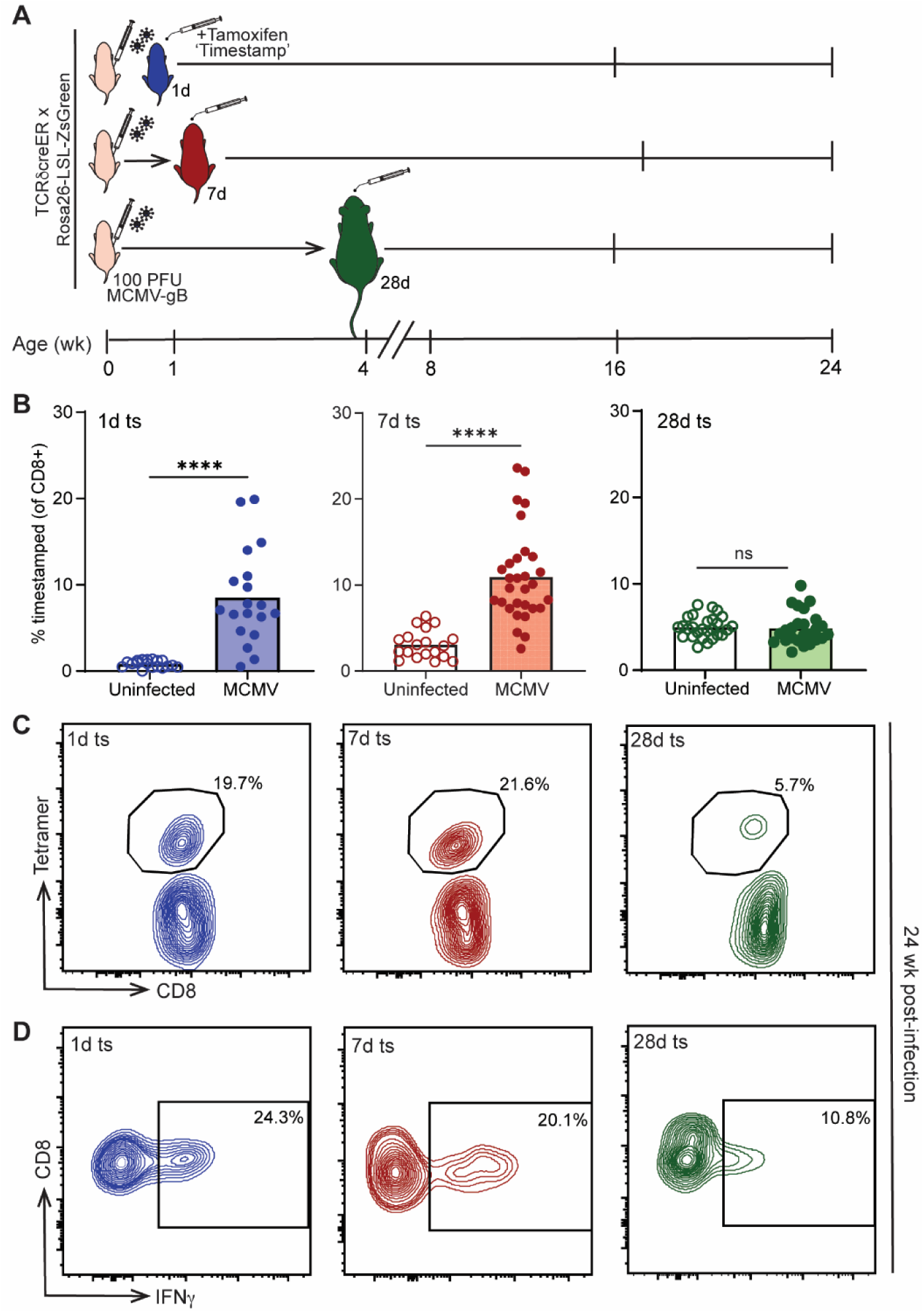
Neonatal CMV infection recruits and maintains CD8+ T cells closest to thymic egress into memory inflation response. (A) Experimental schematic. Newborn mice were infected with MCMV-gB at birth. Uninfected mice were injected with PBS as control. Mice were given tamoxifen at 1 day, 7 days, or 28 days post-birth to ‘timestamp’ CD8+ T cells with a Zsgreen fluorescent tag. (B) Mice were bled at 16-17 weeks post-birth and circulating CD8+ T cells were examined for the percentage marked by flow cytometry (N=6-24 mice). (C) Spleens collected at 21 weeks post-birth were stained for antigen-specific tetramer+ cells. Representative histograms of tetramer+ CD8+ T cells within the timestamped population in the spleen (N = 8 mice per group). (D) CD8+ T cells from the spleen were enriched and gB peptide stimulation with BFA was preformed for 4 hours (N=8 mice per group). Cells were then intracellularly stained for effector molecules. For statistical test of two-groups an unpaired t-test with Mann-Whitney test for correction was performed. Results are shown as mean ± SD or mean only. **p<0.01, ****p<0.0001.

First, we wanted to know if exposure to persistent infection as a neonate expanded the early developmental layers into adulthood. To address this question, we bled the mice at 16-24 weeks post infection and compared the relative numbers of cells at 1d, 7d, and 28d in infected mice to those found in uninfected controls (Fig 2A). Interestingly, the cells made at 1d increased from ∼1% of the bulk CD8+ T cell compartment in uninfected mice to ∼9% in infected mice. The cells made at 7d also increased from ∼3% in uninfected mice to ∼11% in infected mice. In the cells made at 28d, there was no significant difference in the percentage of marked cells between uninfected and infected animals (Fig 2B).

In addition, we asked whether the expansion of cells labelled at 1d and 7d was driven by an accumulation of antigen-specific CD8+ T cells. For this question, we collected spleens at 24 weeks post birth and examined the percentage of 1d, 7d, and 28d marked cells within the tetramer positive population. Among gB-specific CD8+ T cells, ∼20% were composed of 1d marked and another ∼22% were composed of 7d marked cells, while 28d marked cells only constituted ∼6% (Fig 2C, S2A Fig). Thus, CMV-specific cells made closest to the time of infection significantly expand and make up a larger proportion of the total CD8+ T cell compartment.

To confirm whether the expanded 1d and 7d cells retained functionality in adulthood, we restimulated the CD8+ T cells with gB peptide and assessed their ability to produce IFNγ using intracellular flow cytometry. Consistent with an increased proportion of tetramer+ cells, the 1d and 7d marked cells that were restimulated with gB peptide made higher levels of IFNγ (Fig 2D). In contrast, fewer 28d marked cells produced IFNγ, which is likely explained by the lower numbers of gB tetramer positive cells in this population (Fig 2D).

Collectively, our data shows that during neonatal CMV infection, the cells made closest to the time of infection are preferentially recruited into the memory inflation response. These newly made T cells are also known as recent thymic emigrants (RTEs) which are known to possess unique characteristics [30–33]. Strikingly, in our model, these RTEs also persist for long periods of time within the host and retain functionality.

### CD8+ recent thymic emigrants preferentially acquire an effector memory phenotype

We next asked whether the phenotype of CD8+ T cells during memory inflation corresponds to their time of production. We first examined the ‘bulk’ timestamp population and measured T cell differentiation based on their expression of CD44 and CD62L. The vast majority of 1d stamped (95%) and 7d stamped (85%) cells exhibited a more terminally differentiated phenotype (CD44+CD62L-) (S2B-C Fig). In contrast, the 28d stamped CD8+ T cells were comprised of fewer terminally differentiated cells and were the only marked population that contained significant numbers of naïve (CD44-CD62L) and central or virtual memory (CD44+CD62L+) cells.

We next turned our attention to the antigen-specific CD8+ T cell compartment and used CD27 and CX3CR1 to identify different subsets of memory cells within the tetramer+ CD8+ T cell pool. Memory T cells with a central memory phenotype (CD27+ CX3CR1-) can travel between circulation and lymphatics, while more terminally differentiated peripheral (CD27+ CX3CR1+) and effector (CD27-CX3CR1+) memory cells circulate in peripheral tissues [14, 27]. Interestingly, the antigen-specific 1d and 7d marked cells preferentially gave rise to effector memory cells, whereas the 28d marked cells were biased towards forming central memory cells (Fig 3A-B). We also obtained similar results by phenotyping the cells producing IFNγ upon restimulation with peptide. The IFN-producing cells in the 1d and 7d marked population exhibited more of an effector memory phenotype, whereas those made at 28d exhibited more of a central memory cells (S3A-B Fig).

**Figure 3.**
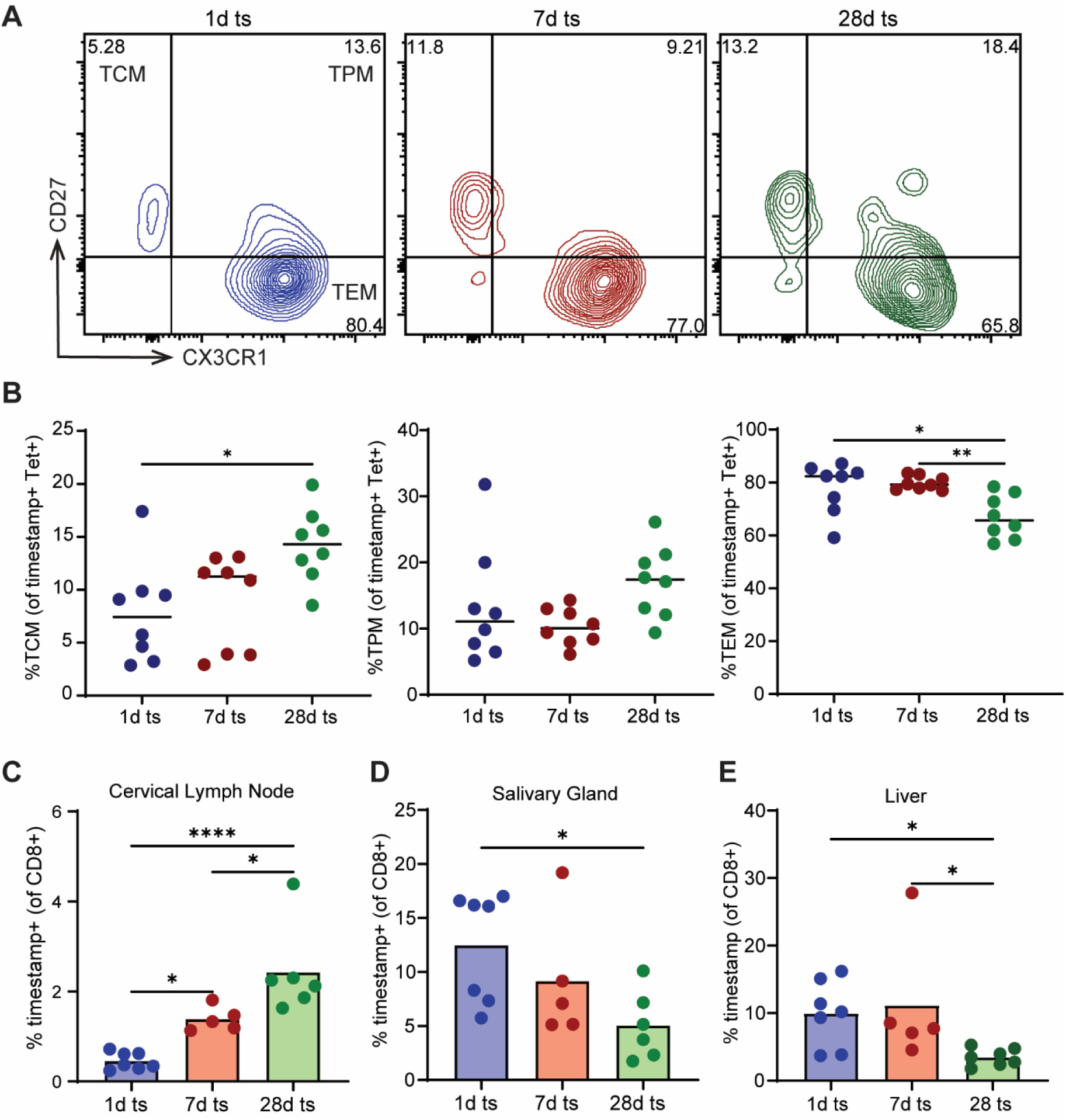
Age of the CD8+ T cell recruited into the memory inflation response dictates phenotype. Newborn mice were infected with MCMV-gB at birth. Uninfected mice were injected with PBS as control. Mice were given tamoxifen at 1 day, 7 days, or 28 days post-birth to ‘timestamp’ CD8+ T cells with a Zsgreen fluorescent tag. Spleens were collected at 21 weeks post-birth and CD8+ T cells were phenotyped by flow cytometry. (A) Representative density plot of CD27 vs CX3CR1 within the tetramer+ sub gates where memory CD8+ T cells were identified as Central Memory (TCM, CD27+ CX3CR1-), Peripheral Memory (TPM, CD27+ CX3CR1+) or Effector Memory (TEM, CD27-CX3CR1+). (B) Quantification of TCM, TPM and TEM phenotype of 1d, 7d and 28d timestamped mice within the tetramer+ population (N=8 mice). (C) Quantification of the percentage of timestamp CD8+ T cells within different tissues. One-way ANOVA with Tukey’s multiple comparisons test was performed. Results are shown as mean. *p<0.05, **p<0.01.

Given that 1d, 7d, and 28d marked cells exhibit a different phenotype, we next asked whether they preferentially localize at different anatomical sites. Compared to 1d marked cells, the 28d marked cells were more significantly enriched in lymphoid tissue (e.g., cervical lymph node) and less abundant in peripheral organs (e.g., salivary gland and liver) (Fig 3C-E). The 7d marked cells exhibited a localization pattern somewhere in between the 1d and 28d marked cells; there were more 7d marked cells in the cervical lymph node compared to 1d marked cells, but there were also more 7d marked cells in the liver compared to 28d marked cells. Together, this data suggests that RTEs have an enhanced capacity to form effector memory cells that migrate to peripheral organs, whereas cells made during the latent stage of infection are biased towards becoming central memory cells and localize in the lymphoid tissue.

### Adult RTEs are preferentially recruited into the inflating pool in CMV infections that occur later in life

We next wondered whether preferential recruitment of RTEs into the CMV response is due to the fact these cells were RTE at the time of infection, or because these cells had a neonatal phenotype? It is possible that after neonatal infection the adult phenotype CD8+ T cells labeled on d28 do not get recruited into the response because neonatal cells are more responsive, because RTE are more responsive, or simply because there is less antigen present at the time adult cells are produced. To test this, we infected mice as adults with MCMV-gB and measured the memory inflation response (Fig 4A). Similar to our neonatal MCMV infection model, we confirmed that memory inflation occurred and leveled off at ∼25% by 8 weeks post infection (Fig 4B, Fig 1B). To look at the recruitment of recent thymic emigrants into the memory inflation response during adult MCMV infection, mice were timestamped at 1d, 7d, and 28d post birth and infected with 1×10^5^ PFU of MCMV-gB at 8 weeks post birth (Fig 4C). In this scenario, the 28d marked cells are now the most recently produced population of CD8+ T cells (i.e., RTEs) at the time of infection.

**Figure 4.**
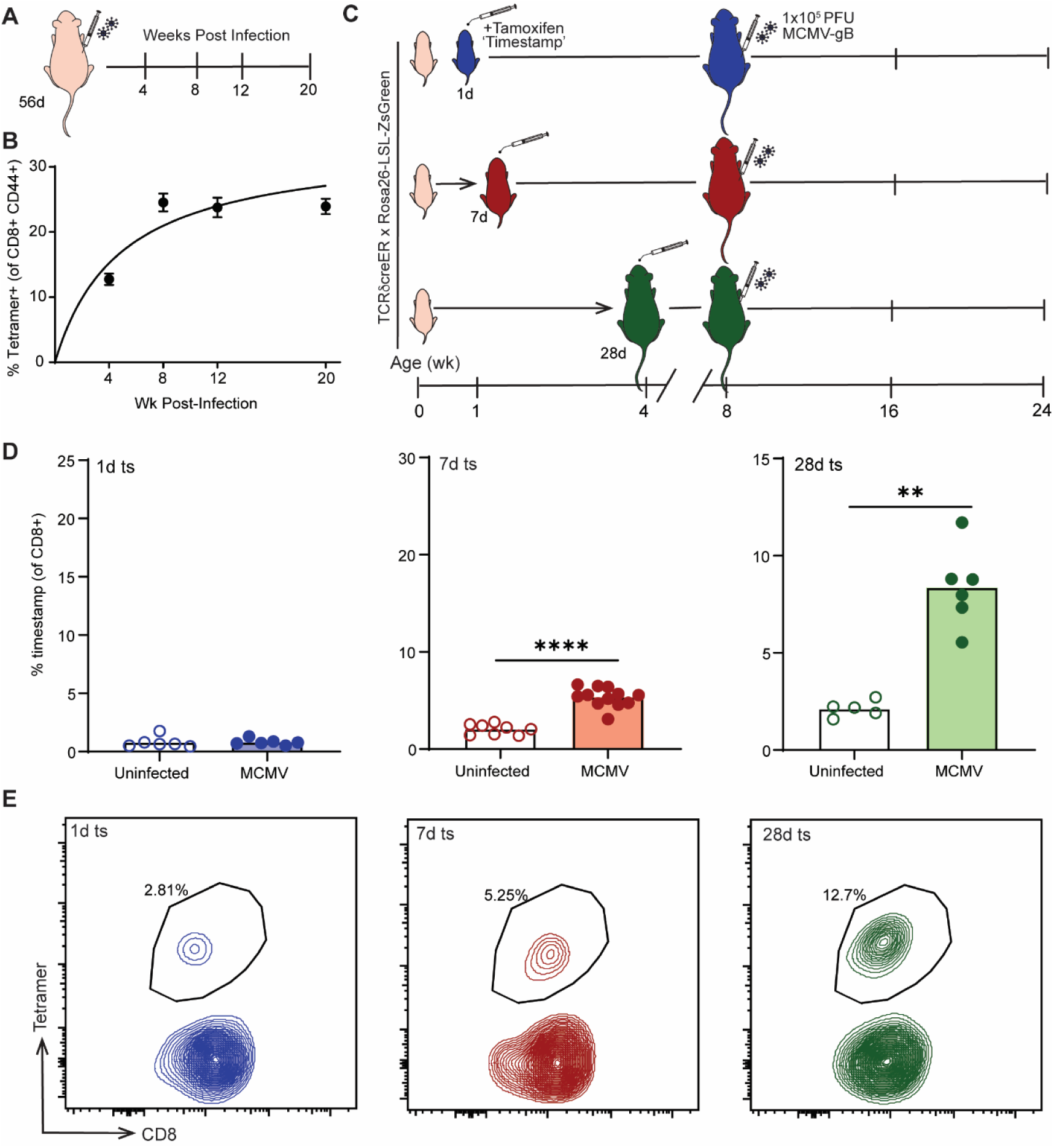
Adult infection with CMV recruits and maintains CD8+ T cells closest to the time of thymic egress into the memory inflation response. (A) Adult mice were infected with MCMV-gB at 56 days post-birth and bled 4, 8, 12 and 20 weeks post-infection. (B) Quantification of CD8+ T cells that were tetramer+. Adult mice were infected with MCMV-gB at 56 days post-birth. Uninfected mice were injected with PBS as control. Mice were given tamoxifen at 1 day, 7 days, or 28 days post-birth to ‘timestamp’ CD8+ T cells with a Zsgreen fluorescent tag. Mice were bled at 16 weeks post-birth and circulating Zsgreen+ CD8+ T cells were examined by flow cytometry. (C) Experimental schematic. (D) The percentage of CD8+ T cells that are Zsgreen+ tetramer positive (N = 5-12 mice). (E) Spleens were collected from adults at >24 weeks post-birth and CD8+ T cells were stained for gB tetramer. Representative scatter plot of tetramer+ CD8+ T cells within the timestamped population. For statistical test of two-groups an unpaired t-test with Mann-Whitney test for correction was performed. Results are shown as mean. *p<0.05, ***p<0.001, ****p<0.0001

To control for the age of the host, we bled the mice at 16 weeks post birth to match the timepoint examined after neonatal infection and measured the percentage of cells marked at different ages. Interestingly, there was no significant difference in the percentage of total 1d stamped cells between uninfected and adult MCMV infected animals (Fig 4D). However, there was a significant increase in the percentage of 7d and 28d marked cells in adult MCMV infected animals, with the 28d being the most pronounced population (Fig 4D). We then examined the differentiation and activation status of the marked cells and observed less differentiated (CD44+CD62L+) cells in the 1d and 7d marked populations and more terminally differentiated (CD44+CD62L-) cells in the 28d marked population (S4A-B Fig).

We next collected spleens at 24 weeks of age and measured the antigen-specific tetramer response in the adult-infected mice. The 28d marked cells had a significant increase in the percentage of tetramer positive cells compared to both 1d and 7d marked cells (S4C Fig). In fact, only ∼3% of the 1d marked cells were tetramer positive, while ∼5-6% of the 7d and ∼13% of the 28d marked cells contributed to the antigen response (Fig 4E). Thus, the preferential recruitment of CD8+ T cells produced closest to the time of infection and their bias towards becoming effector memory is not a unique attribute of neonatal infections, but instead appears to be a common feature of the CD8+ T cell response to CMV infection in later life.

### Post-thymic maturation is a key determining factor of T cell recruitment during CMV infections

Up to this point, we have shown that RTEs are preferentially recruited into the memory inflation pool in both neonatal and adult infections. While these experiments provide important information on the CD8+ T cell response to infection at different stages of life, there are several key variables not controlled for in each infection scenario. For example, in the adult infected mice, it is possible that the 1d marked cells are not efficiently recruited into the response not because they have undergone more post-thymic maturation, but because they are simply outcompeted by the 28d marked cells, which are derived from a different progenitor and likely express more diverse TCRs [22, 29, 34, 35]. To test this, we designed an experiment to control for the age of the cells and compare how RTEs of different developmental origins respond to infection in the same environment.

To differentiate between these possibilities, we performed a thymic transplant experiment where the thymus from a fetal (TdTomato) timestamp donor mouse was engrafted under the kidney capsule of an adult (ZsGreen) timestamp mouse (Fig 5A). Thus, when tamoxifen is administered, it will mark CD8+ T cells of fetal (red) and adult (green) origin at the same time and allow us to directly compare fetal RTEs vs adult RTEs in the same (adult) environment. If developmental origin is a key determinant, then we would expect to see either the fetal or adult RTE population get preferentially recruited into the adult response. On the other hand, if post-thymic maturation is the major factor, then both fetal and adult cells are RTE so should behave similarly. On day 5 post thymic transplant, the recipient mice were infected with MCMV-gB and bled every 4 weeks to measure expansion of the marked fetal and adult cells (Fig 5A). We found that there was no statistical difference between the expansion of both fetal and adult CD8+ T cells at 4, 8 or 12 weeks post infection.

**Figure 5.**
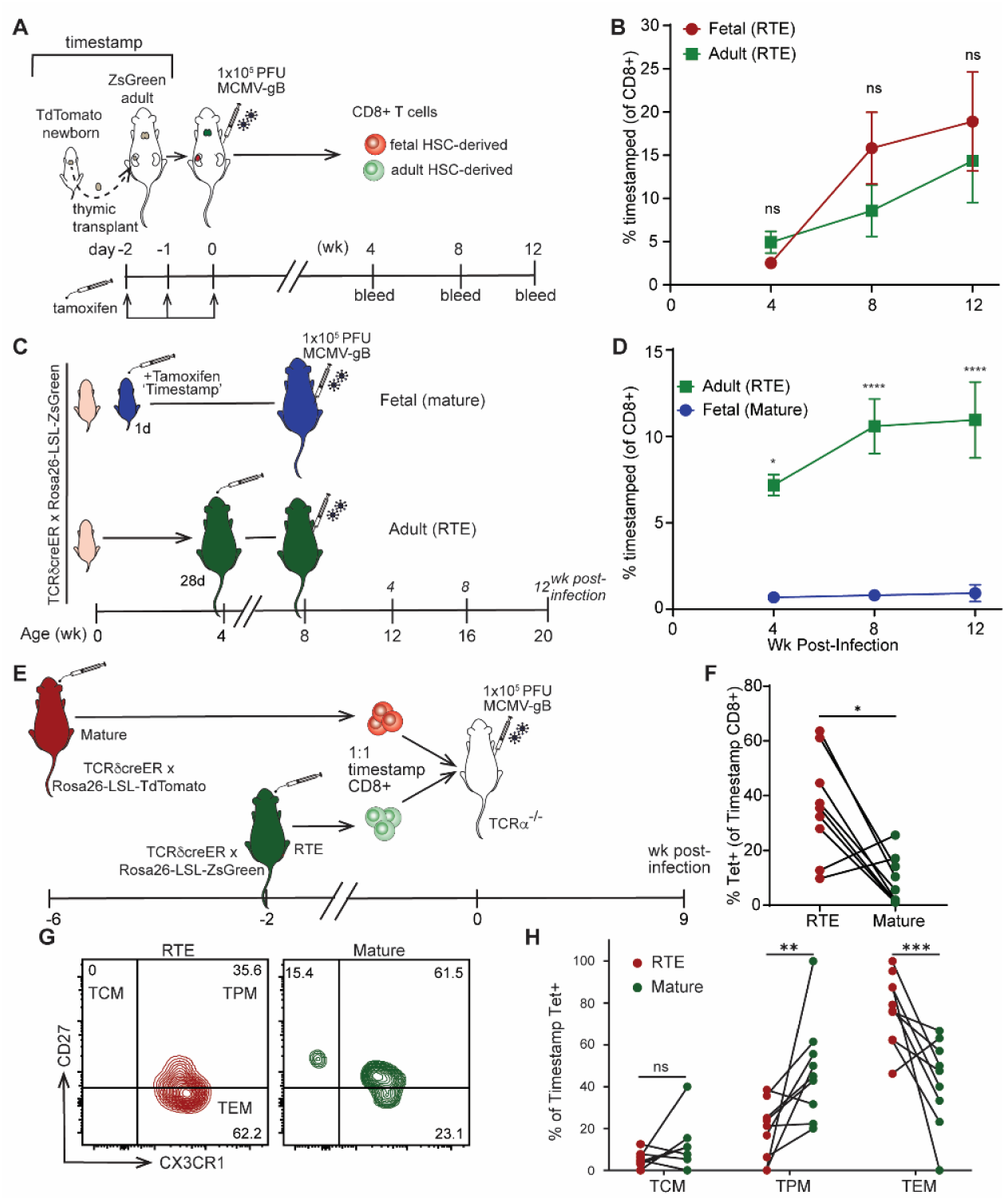
Recent thymic emigrants are preferentially recruited into the memory response, maintained, and are terminally differentiated. (A) Newborn thymuses were collected from Ai9+ timestamp reporter mice and surgically transplanted under the kidney capsule into adult (>8 week) Zsgreen+ timestamp reporter mice. Mice were administered tamoxifen for 3 days and then infected with MCMV-gB two days after marking. At 4, 8 and 12 weeks post-infection, mice were bled and CD8+ T cell memory response was measured by flow cytometry. (B) The percentage of CD8+ cells that were Zsgreen/Ai9 marked cells (N=16-21 mice). (C) Mice were administered tamoxifen at 1d to mark ‘fetal’ layer or 28d post-birth to mark ‘adult’ layer. All mice were infected at 56 days post-birth and bled 4, 8 and 12 weeks post-infection. (D) The percentage of CD8+ T cells that were from 1d or 28d marked layers. E) Adult Zsgreen+ mice were marked with tamoxifen 6 weeks prior to harvest, while adult Ai9+ mice were marked with tamoxifen 2 weeks prior to harvest. CD8+ T cells were isolated from the spleen. A 1:1 ratio of Zsgreen+ and Ai9+ CD8+ T cells were pooled and co-transferred into adult TCRa^−/−^ animals. The next day recipient animals were infected with MCMV-gB. Mice were bled at 9 weeks post-infection and CD8+ T cell response was measured by flow cytometry. F) The percentage of Zsgreen/Ai9 marked CD8+ T cells that were Tetramer+ (N=9). G) Representative 2-way FACS plot of CD27 vs CX3CR1. H) The percentage of tetramer+ timestamped CD8+ T cells that adopted a central memory (TCM), peripheral memory (TPM) or effector memory phenotype (TEM). A paired t-test with Wilcoxon matched pairs signed rank test performed for correction. Results are shown as pair-wise comparison where each connected point is one mouse. *p<0.05, ***p<0.001, ****p<0.0001.

To validate the significance of our thymic transplant findings, we also used our previous model in Figure 4, and examined the fetal or adult layer of CD8+ T cells in mice infected as adult with MCMV-gB at the same timepoints as those used in the transplant studies (Fig 5C). In this scenario, all fetal marked cells would be “mature”, while the adult marked cells would be closest to RTE population (Fig 5C). As expected, only the adult RTE cells expanded at 4, 8, and 12 weeks post infection, while the fetal mature cells failed to expand during the course of infection (Fig 5D). Taken together, this data shows that developmental origin doesn’t dictate the response to memory inflation, but instead suggests that the age of the cell is a key determining factor.

Another variable that was not controlled for in earlier experiments was the precursor number. For example, it is possible that we see more RTEs respond to adult infection because there are more 28d marked cells than 1d marked cells at the time of infection. Thus, we felt it was important to design an experiment to directly compare equal numbers of adult (28d marked) cells in the same environment that only differ in maturation (RTEs vs mature cells). To address this question, we again turned to our timestamp model. In this case, ZsGreen+ timestamp mice were given tamoxifen at 6 weeks prior to infection, making the marked CD8+ T cells mature at the time of harvest. A second group of adult TdTomato+ timestamp mice were given tamoxifen at 28 days of age, and infected two weeks later (marked CD8+ T cells were RTE). Marked cells from both donors were harvested and combined in a 1:1 ratio and transferred into TCRα^−/−^ animals that were then infected the following day (Fig 5E). We found that the TdTomato+ (RTE) population was more readily recruited into the antigen-specific tetramer response than the mature ZsGreen+ (mature) population, starting at 2 weeks post infection (Fig 5F). Consistent with our previous observations, RTEs that were recruited into the memory inflation response more readily adopted a terminally differentiated effector memory phenotype (Fig 5G-H). Altogether, our findings indicate that RTEs are a major source of cells recruited into the memory inflating pool that are retained for the life of the animal.

## Discussion

Previous work has suggested that CD8+ T cell responses are maintained over the course of persistent infection because they are replenished by a constant pool of short-lived effector cells [4, 36, 37]. Conceptually, one could hypothesize that the short-lived effectors replacing the pool are recruited from new RTEs that are constantly produced during latent infection. However, as we demonstrate, this is not the case. Specifically, we did not observe an equal recruitment of CD8+ T cells made at different stages of life or ongoing recruitment of cells during persistent infection. Instead, the CMV-specific memory pool is maintained by cells produced closest to when the host was first infected. This leads us to propose that there is a ‘critical window’ for CD8+ T cells to be recruited into the inflating pool, which corresponds to how recently they were exported from the thymus at the time of infection.

Although the underlying basis for why RTEs are preferentially recruited into the CD8+ T cell response after persistent viral infection is not clear, several possibilities are worth considering. First, RTEs are more biased towards becoming short-lived effectors than mature CD8+ T cells, even when responding to infection in the same host [31, 32]. The enhanced propensity for RTEs to undergo effector cell differentiation may allow them to more rapidly fill the pool that is available to sustain memory inflation. Second, RTEs express higher levels of VLA-4 than mature cells, which enhances their ability to migrate into various tissues [30]. The increased ability of RTEs to gain access to peripheral tissues may enable them to encounter more CMV antigens than mature cells and undergo memory inflation. Third, RTEs have a superior ability to respond to low-affinity antigens [30]. Thus, it is possible that RTEs are preferentially recruited into the inflating response because they can respond to CMV antigens that are not recognized by mature cells. Future experiments are required to examine the relative contribution of these possibilities.

Our work sheds new light on the competing models of memory inflation. Work from the Oxenius lab suggested that KLRG1-TCF1+ central memory-like CD8+ T cells maintain the inflationary pool of T cells during CMV [37, 38]. Other evidence from the Klenerman lab suggested that a self-proliferating pool of peripheral memory CD8+ T cells replenish the inflationary pool after CMV infection [14]. However, Gerlach et al. demonstrated that peripheral memory CD8+ T cells are highly plastic and exist in a transient state where they can revert back to a central memory phenotype or terminally differentiate into effector memory phenotype, which could explain why there are currently multiple models of how the inflationary pool is maintained [27]. Importantly, our studies do not need to be viewed as a new or opposing model. For example, our results support the prevailing idea that long-lived populations maintain effectors. However, we show that this population is selectively recruited from RTEs present at the time of infection (rather than from the total CD8+ T cell pool).

One of the most interesting findings of our study is that the timing of infection dictates the types of CD8+ T cells that are maintained in the memory pool. Whereas infections in early life expand the neonatal layer of CD8+ T cells, infections in adulthood promote an increase in adult CD8+ T cells. Thus, an important question is, are there are consequences to having a larger number of neonatal cells in the adult memory pool? Previous work has indicated that CMV infection contributes to a large amount of immune variation in the human population [25, 39, 40]. Whether the timing of infection explains some of the variability in outcomes among CMV patients remains an open question.

Lastly, our work provides new insight into age-related differences in the CD8+ T cell response to persistent viral infections. Neonates and lymphoreplete individuals have a larger and more robust pool of RTEs within their CD8+ T cell compartment [41, 42]. It is interesting to speculate whether memory inflation occurs at a more rapid rate because of this larger RTE pool. Humans seropositive for CMV have a diverse percentage of CMV-specific CD8+ T cells. In one study, seropositive individuals ranged from <2% to >40% (average of 10.2%) of their memory CD8+ T cell pool specific to HCMV [8]. It will be important to determine whether the size of the RTE pool at the time of infection has a direct impact on the rate, size, and phenotype of the inflationary pool within individuals. This may have particular implications for individuals acquiring CMV after HSCT or other lymphodepleting therapy, which may affect RTE frequency at the time of infection. Our data showing that RTEs are preferentially recruited into the response means that the age of the cell present at the time of infection will need to be taken into account when developing future strategies to boost immunity against persistent viral infections at difference stages of life.

## Materials and Methods

### Animals

C57BL/6NCR mice were purchased from Charles River. gBT-I TCR transgenic mice (specific for the HSV-1 glycoprotein gB498–505 peptide, SSIEFARL) were provided by Dr. Janko Nikolich-Zugich (University of Arizona, Tucson, AZ). ZsGreen, Ai9, TCRδCre-ERT2 and TCRα^−/−^ mice were purchased from Jackson Laboratories. Male and female mice were used for all experiments unless otherwise specified. All results are pooled experiments from different batches of mice. All experiments in this study were conducted in accordance with the recommendations in the Guide for the Care and Use of Laboratory Animals of the National Institutes of Health and protocols reviewed and approved by the Institutional Animal Care and Use Committee at Cornell University.

### Timestamping Mice

Mice were generated by breeding TCRδCre-ERT2 with ZsGreen or Ai9 reporter mice. To mark 1d (fetal) T cells, 2.5 mg of tamoxifen was administered to dams by oral gavage, 3 times in 12 hour increments. Pups received the tamoxifen through lactation. To mark 7d (neonatal) T cells, pups were administered tamoxifen 0.25 mg by oral gavage. To mark 28d (adult) T cells mice were given 2.5 mg of tamoxifen by oral gavage.

### MCMV Infection

Recombinant MCMV expressing the MHC class I-restricted CTL epitope HSV gB498–505 (SSIEFARL), designated in the text as MCMV-gB, was provided by Dr. Cicin-Sain (Helmholtz Centre for Infection Research, Germany). Newborn pups (<24 hours postpartum) were infected intraperitoneally (i.p.) with 100 plaque forming units (PFU) for the newborn MCMV group of animals. Adult mice were infected i.p. with 1×10^5^ PFU for adult MCMV group of animals.

### Flow Cytometry

Mouse antibodies for CD4 (Gk1.5), CD8a (53-6.7), CD27 (LG.3A10), CD44 (IM7), CD49d (R1-2), CD62L (MEL-14), CD103 (M290), CD122 (TM-B1), CX3CR1 (SA011F11), TCRβ (H57-597), TCRγδ (eBioGL3), Perforin (S16009A), TNFα (MP6-XT22), IFNγ (XMG1.2), Granzyme B (GB11), were purchased from BD Bioscience, BioLegend or ThermoFisher Scientific. Fixable viability dye eFluor 780 was purchased from TheremoFisher Scientific. Biotinylated monomers against gB peptides were obtained from the National Institutes of Health Tetramer Core Facility and tetramerized with streptavidin-linked fluorophores within our lab.

Whole blood was lysed for red blood cells by addition of dH20 followed by quickly adding 4x PBS. Lysis was performed twice and then washed twice with MACS solution (1x PBS, 0.5% BSA, 2mM EDTA). Spleens were homogenized using a syringe over a 40 uM filter. CD8+ T cells were bead enriched from the spleens by positive selection with CD8a microbeads (Miltenyi). Spins between each was and resuspension was at 500g for 3 minutes in 96 well plates. Cells were stained in antibody cocktail suspended in Brilliant Stain Buffer (BD Biosciences) and MACS. Cells were stained for 30 minutes without tetramer or 60 minutes with tetramer at 4°C in the dark. For surface staining cells were fixed with IC fixation kit (Invitrogen). For intracellular staining cells were permeabilized using Foxp3/Transcription Factor Staining Buffer (Invitrogen). Cells were incubated with intracellular antibodies in Brilliant Stain Buffer (BD Biosciences) and MACS for 30 minutes at 4°C in the dark. All cells were resuspended as a final single cell suspension in MACS.

For gating schemes lymphocytes were identified by FSC/SSC. Singlets were gated on FSC/FSH and SSC/SSH. Conventional CD8+ T cells were identified by TCRγδ-/CD4-/TCRβ+/CD8a+. Antigen-specific cells were identified by tetramer staining, timestamp cells were ZsGreen+, true naïve CD8+ T cells were gated as CD49d-/CD44-/CD122-, virtual memory CD8+ T cells were gated as CD49d-/CD44+/CD122+ and true memory (TMEM) CD8+ T cells were gated as (CD49d+/CD44+). Within the TMEM population central memory (TCM) was identified as CD27+CX3CR1-, peripheral memory (TPM) as CD27+ CX3CR1+ and effector memory (TEM) as CD27-CX3CR1+.

All experiments were run on BD FACSymphony A3, BD FACSymphony A5 or Accuri C6. Data was then analyzed on FlowJo Version 10.

### In vitro stimulation

CD8+ T cells from whole blood or spleen were stimulated with 1 × 10^−7^ M of SSIEFARL peptide with 1.5 µg/ml Brefeldin A (Millipore Sigma) for 4 hours at 37°C resuspended in RP-10 media (RPMI 1640, 10% fetal bovine serum, 1x L-glutamine, 1x penicillin/streptomycin).

### Cell Counts

Zsgreen reporter mice were bred for cell enumeration. Mice were either left uninfected or infected with MCMV at D0 or D56 same as experimental groups. Spleens, cervical lymph nodes and thymus were collected and processed into a single cell suspension. Total cell counts were enumerated by a MoxiZ (ORFLO) and the absolute number of CD8+ T cells were back calculated by the percentage of CD3+ CD8a+ cells on flow cytometry.

### Co-transfer Experiment

ZsGreen timestamp mice at 6 weeks of age were marked with tamoxifen for 3 days in 24-hour increments, 6 weeks prior to harvesting the spleen. TdTomato timestamp mice at 4 weeks of age were marked with tamoxifen for 3 days in 24-hour increments, 2 weeks prior to harvesting the spleen. CD8+ T cells were isolated from Zsgreen+ and TdTomato+ spleens at the same time. Cells were enumerated and a 1:1 of 200k Zsgreen+ and 200k TdTomato+ CD8+ T cells were pooled and transferred into TCRα^−/−^ animals by intravenous injection. The next day TCRα^−/−^ recipients were infected with MCMV-gB and subsequently retro-orbitally bled 9 weeks post-infection.

### Thymic Transplants

Transplants were conducted as previously described with modification [22]. Thymus lobes were isolated from 1d TdTomato+ timestamp mice. The lobes were placed surgically under the kidney capsule of a 7 week old Zsgreen+ timestamp recipient mouse. To mark the thymocytes, 5 mg of tamoxifen was administered by oral gavage for 3 days post-surgery in 24 hour increments. Mice were infected with MCMV-gB at 5 days post-surgery and bled retro-orbitally at 4, 8 and 12 weeks post-infection.

### Statistical Analysis

All error bars are represented as mean plus or minus standard deviation or standard error of the mean. For comparison between two groups an Unpaired t test was used; if the standard deviation was not equal a Welch’s correction was applied. All statistics were performed in GraphPad Prism Version 9. For comparison between three or more groups an One-way or Two-way ANOVA was used for appropriate test correction (Bonferroni, Tukey) as indicated in each figure legend. The regression line shown in Figure 1A, Figure 4A, and Supplemental Figure 1A was an One phase exponential association as defined by Y=Ymax*(1-exp(-K*X)) where the line starts at zero and ascends to Ymax with a rate constant K, half time is 0.69/K.

## Acknowledgements

We thank the Cornell Center for Animal Resource and Education (CARE) for expert mouse breeding assistance. Cell sorting was done at Cornell University’s Flow Cytometry Facility in the Biotechnology Resource Center (RRID:SCR_021740). This work was supported by National Institute of Health Grants R01AI105265, R01HD107798, R01AI110613, and R21AI142382 (to B.D.R). Trainee (Z.T.H.) support was provided by the American Association of Immunologist Intersect Fellowship for Computational Scientists and Immunologist. M.P.D is funded by an NHMRC (Australia) Investigator grant #1173027.

