## Supplemental Figures for "Recent thymic emigrants are preferentially recruited into the memory pool during persistent infection"

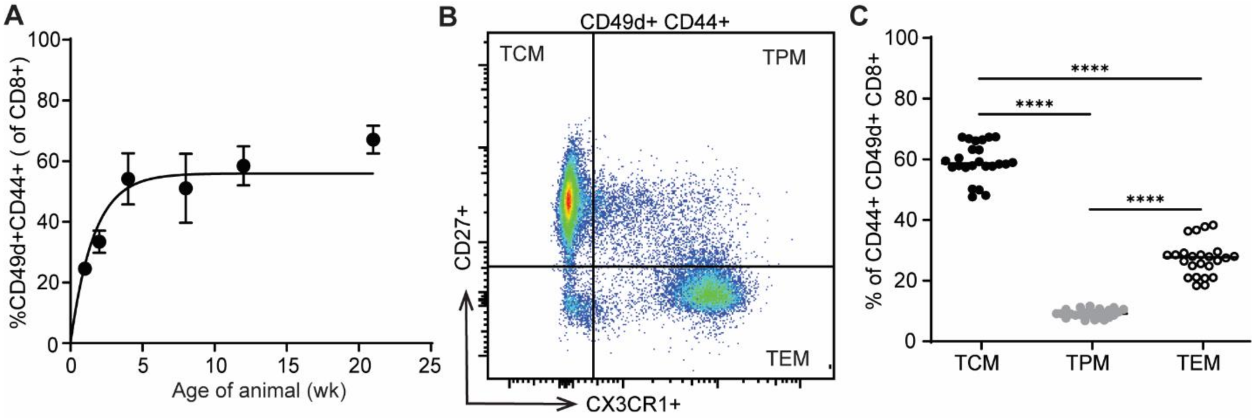


**Supplemental Figure 1. Neonatal infection with MCMV undergoes inflation of 'antigen-experienced' CD8+ T cells.** Newborn mice were infected with MCMV-gB at birth. CD8+ T cells were isolated from the spleen at 1, 2, 4, 8, 12 and 21 weeks post-birth. (A) CD8+ T cells were stained for CD44 and CD49d to measure 'antigen-experienced' cell by flow cytometry (N = 4-10 mice). (B) CD8+ T cells within the CD44+ CD49d+ sub-gate were stained for CD27 vs CX3CR1 to distinguish memory phenotype (Central Memory [TCM, CD27+ CX3CR1-], Peripheral Memory [TPM, CD27+ CX3CR1+], Effector Memory [TEM, CD27- CX3CR1+]). Representative 2-way FACS plot of CD27 vs CX3CR1. (C) Quantification of TCM, TPM and TEM CD8+ T cells within the CD44+ CD49d+ sub-gate. An ordinary One-way ANOVA with Tukey's multiple comparisons test was performed. Results are shown as mean ± SD or mean only. ***$p<0.001$, ****$p<0.0001$.

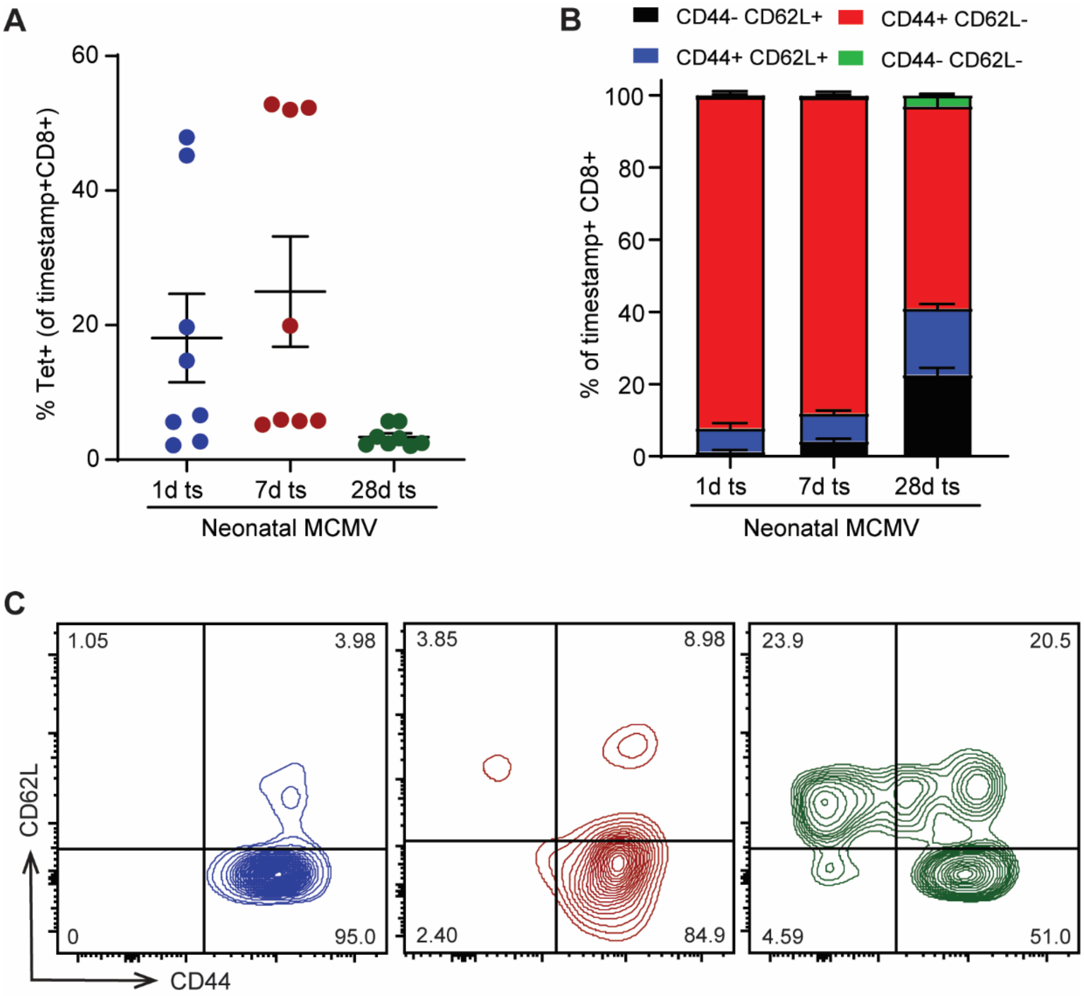


**Supplemental Figure 2. CD8+ T cells made closest to neonatal infection (RTEs) are preferentially recruit into the tetramer response.** Newborn mice were infected with MCMV-gB at birth. Uninfected mice were injected with PBS as control. Mice were given tamoxifen at 1 day, 7 days, or 28 days post-birth to ‘timestamp’ CD8+ T cells with a Zsgreen fluorescent tag. Spleens were collected at 21 weeks post-birth. (A) Quantification of the percentage of timestamp CD8+ T cells that were tetramer+ (N = 8 mice per group). (B) Percentage of total timestamp CD8+ T cells that adopted an CD44 vs CD62L phenotype. (C) Representative 2-way plot of CD44 vs CD62L on total timestamp CD8+ T cells. An ordinary One-way ANOVA with Tukey's multiple comparisons test was performed. Results are shown as mean ± SEM.

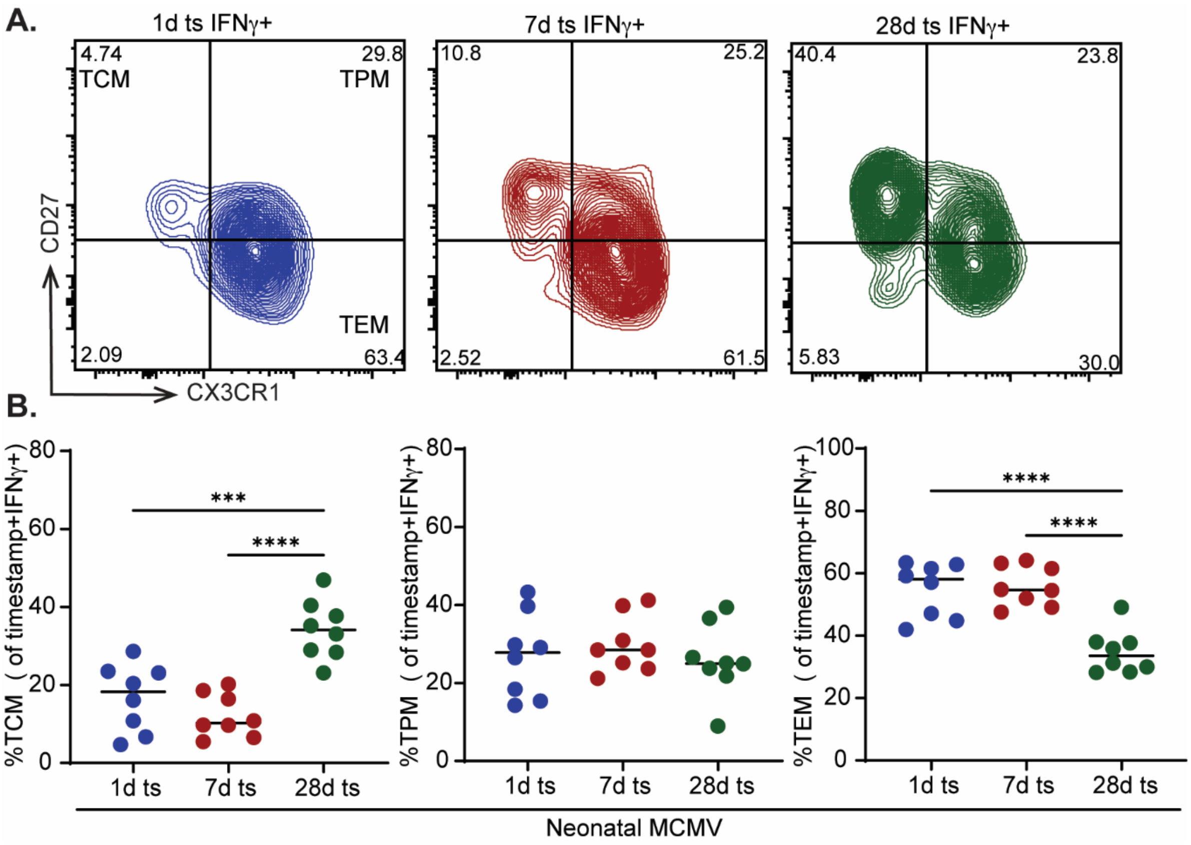


**Supplemental Figure 3. Antigen-specific CD8+ T cells made closest to neonatal MCMV infection (RTEs) adopt a more terminally differentiated phenotype.** Newborn mice were infected with MCMV-gB at birth. Uninfected mice were injected with PBS as control. Mice were given tamoxifen at 1 day, 7 days, or 28 days post-birth to 'timestamp' CD8+ T cells with a Zsgreen fluorescent tag. Spleens were collected at 21 weeks post-birth. CD8+ T cells from the spleen were enriched and gB peptide stimulation with BFA was preformed for 4 hours. Cells were then intracellularly stained for effector molecules. (A) Representative 2-way FACS plot of CX3CR1 vs CD27 within IFNy+ CD8+ population (N=8 mice each group). (B) Quantification of percent TCM, TPM or TEM of IFNy+ CD8+ population (N=8 mice each group). An ordinary One-way ANOVA with Tukey's multiple comparisons test was performed. Results are shown as mean ± SD or mean only.

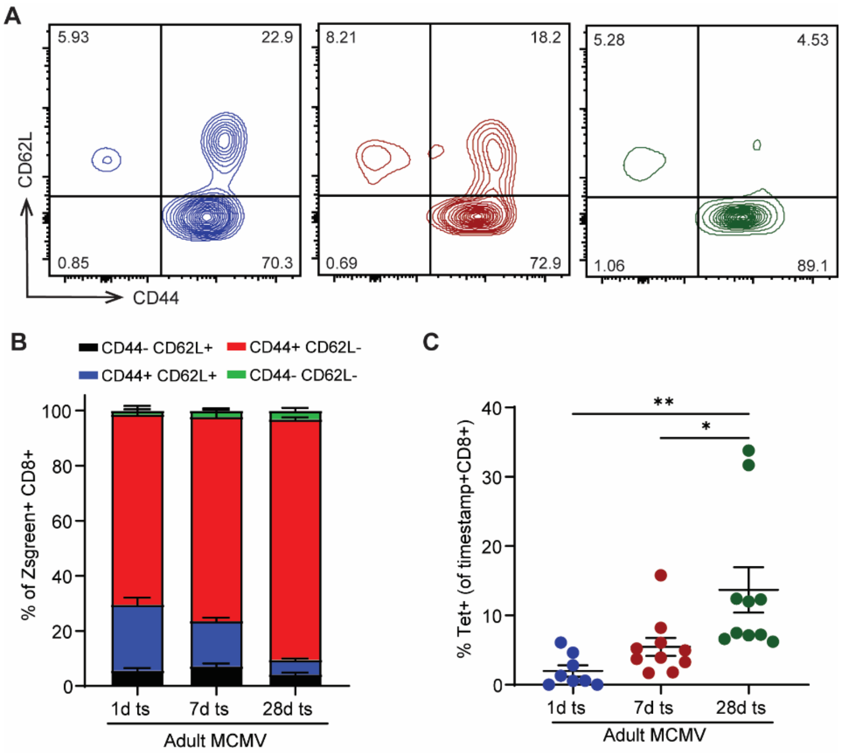


**Supplemental Figure 4. Cells made closest to adult MCMV infection (RTEs) adopt a more terminally differentiated and antigen-experienced phenotype.** Adult mice were infected with MCMV-gB at 56 days post-birth and spleens were collected from adults at >24 weeks post-birth. CD8+ T cells were stained for CD44 vs CD62L to determine differentiation status. (A) Representative 2-way plot of CD44 vs CD62L on total timestamp CD8+ T cells. (B) Quantification of CD44 vs CD62L on total timestamp CD8+ T cells. (C) Quantification of tetramer+ timestamp CD8+ T cells (N=8-10 mice). An ordinary One-way ANOVA with Tukey's multiple comparisons test was performed. Results are shown as mean ± SEM.
